## Supplemental Data 1 for "FunHoP analysis reveals upregulation of mitochondrial genes in prostate cancer"

### Supplementary material

Supplementary Figures S1, S2 and S3

The figures show different aspects of mitochondrial genes in the dataset, using the consensus data on subcellular localization. In each figure the four pathways chosen for further analysis are indicated.

#### Figure S1

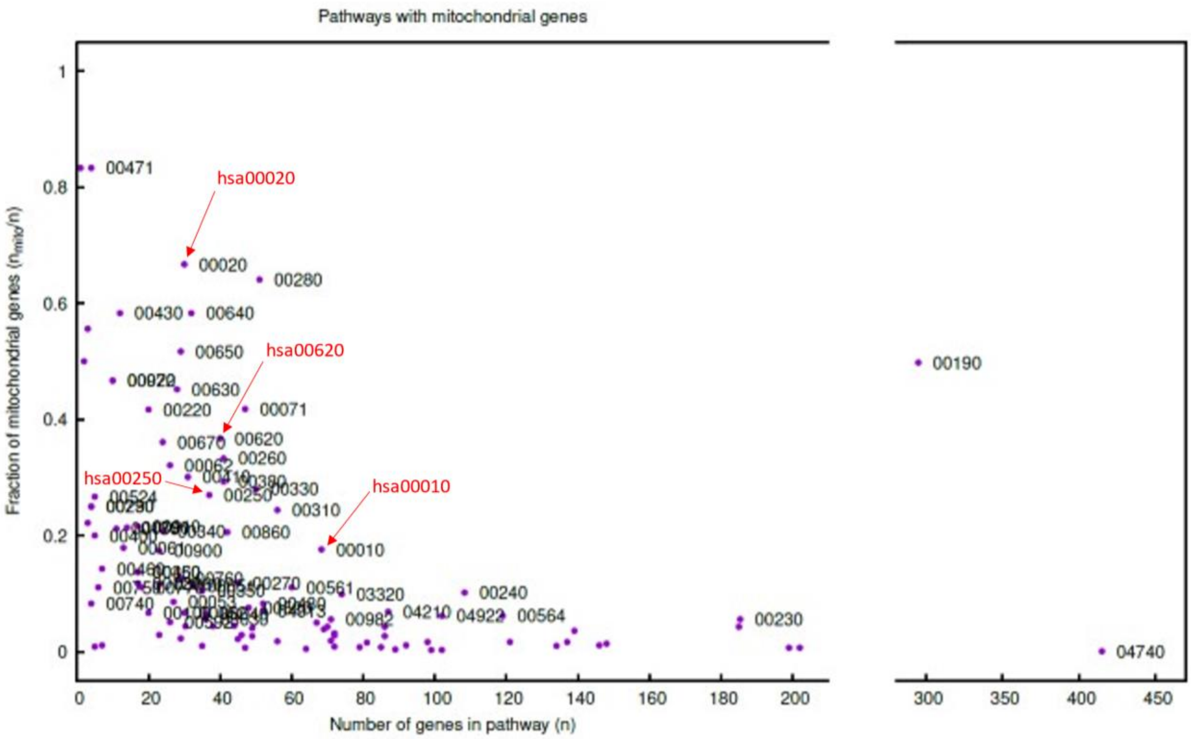


This figure shows the fraction of genes in each pathway that are mitochondrial genes, relative to the number of genes in each pathway. We can for example see that for hsa00280 just over 60% of approximately 50 genes are mitochondrial. The figure shows that the number of pathways with a high percentage of mitochondrial genes is limited.

**Figure S2**

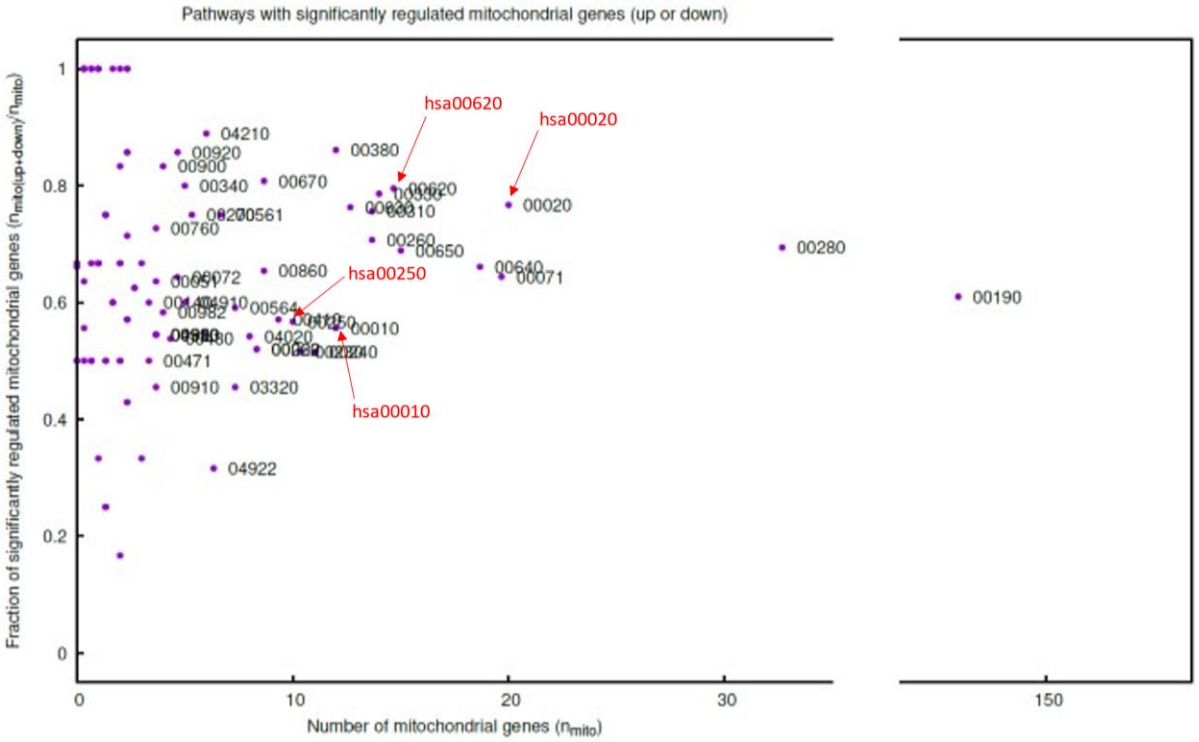


This figure shows the fraction of mitochondrial genes in each pathway that are significantly regulated, both up and down. We see for example that for has00380 more than 80% of the at least 10 mitochondrial genes are significantly regulated. The figure shows clearly that in most pathways where some mitochondrial genes are found, these genes are in general actively regulated.

**Figure S3**

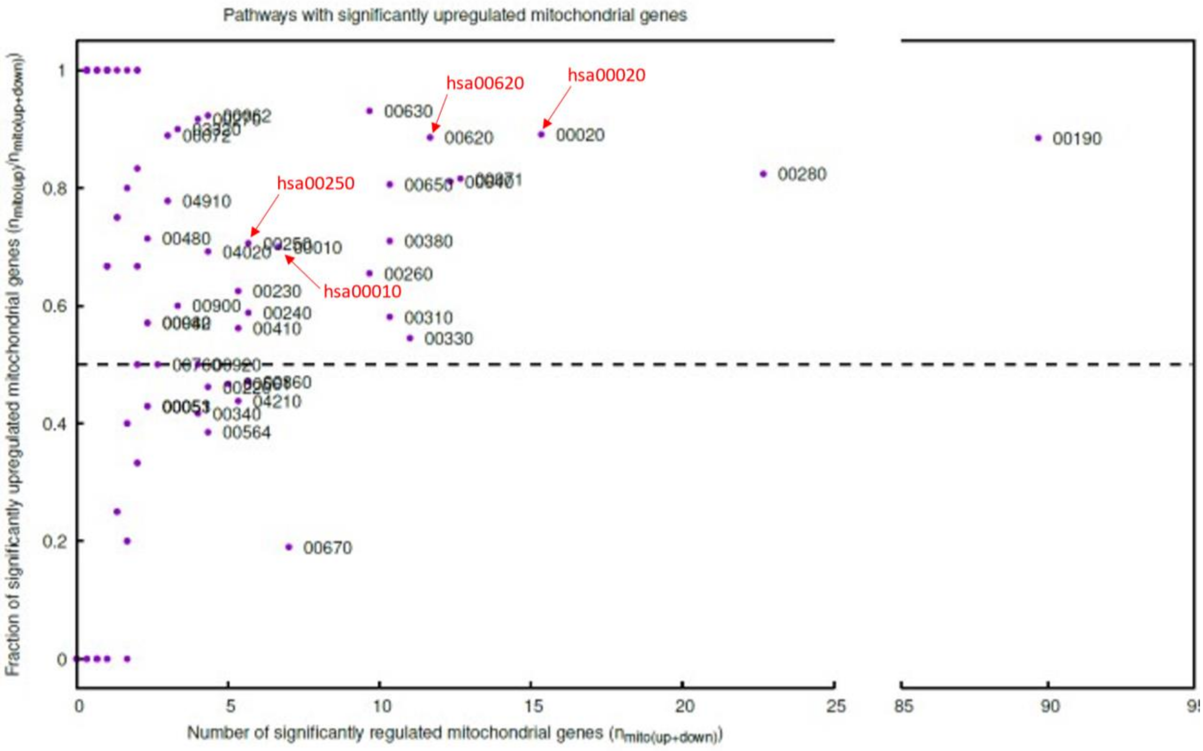


This figure shows the fraction of significant regulated mitochondrial genes that are upregulated. We can for example see that for hsa00020 approximately 90% of the around 15 actively regulated mitochondrial genes are upregulated. The figure shows clearly that upregulation of mitochondrial genes is the most common situation, and that pathways with downregulated mitochondrial genes in general has very few such genes.

Supplementary Tables S1 and S2

**Table S1** – Overlap between categories when comparing the datasets

| *HPA* | *SubCell* | | | | | | |
|---|---|---|---|---|---|---|---|
| | Uncl | Uncrt | Mito | Unkn | Secr | Nucl | Cyto |
| Uncrt | 4 | 5 | 9 | 51 | 58 | 26 | 133 |
| Nucl | 3 | 17 | 7 | 73 | 47 | 282 | 213 |
| Mito | 4 | 8 | 141 | 25 | 15 | 6 | 25 |
| Unkn | 17 | 21 | 60 | 829 | 174 | 44 | 130 |
| Secr | 8 | 16 | 8 | 122 | 215 | 17 | 102 |
| Cyto | 5 | 8 | 8 | 86 | 45 | 15 | 259 |
| *HPA* | *BUSCA* | | | | | | |
| | Memb | Cyto | Unkn | Mito | Nucl | Extra | |
| Uncrt | 67 | 130 | 0 | 24 | 39 | 26 | |
| Nucl | 66 | 251 | 2 | 41 | 265 | 17 | |
| Mito | 27 | 32 | 1 | 154 | 4 | 6 | |
| Unkn | 739 | 184 | 46 | 101 | 60 | 145 | |
| Secr | 231 | 111 | 2 | 23 | 37 | 84 | |
| Cyto | 60 | 244 | 6 | 33 | 56 | 27 | |
| *BUSCA* | *SubCell* | | | | | | |
| | Uncl | Uncrt | Mito | Unkn | Secr | Nucl | Cyto |
| Memb | 16 | 14 | 20 | 726 | 334 | 25 | 55 |
| Cyto | 6 | 24 | 19 | 181 | 79 | 105 | 538 |
| Unkn | 0 | 2 | 12 | 27 | 5 | 3 | 8 |
| Mito | 4 | 15 | 172 | 62 | 24 | 19 | 80 |
| Nucl | 5 | 11 | 4 | 65 | 22 | 213 | 141 |
| Extra | 10 | 9 | 6 | 125 | 90 | 25 | 40 |

Mito – Mitochondria; Nucl – Nuclear / Nucleus; Cyto – Cytosol / Cytoplasm; Memb – Membrane; Extra – Extracellular; Secr – Secretory; Unkn – Unknown; Uncl – Unclassified; Uncrt - Uncertain

This table shows the overlap as number of genes between different categories when comparing HPA data to SubCell, HPA to BUSCA, and BUSCA to SubCell.

For making Figure 1 the BUSCA categories Extracellular and Membrane were combined as Secretory. The data here seems to support this, as these two categories show the largest overlap to Secretory both for HPA and SubCell (although most of the Extracellular and Membrane genes are Unknown (i.e., without classification) in the experimental data).

**Table S2** – Comparison of predicted and experimental localization data

| | TP | FP | FN | TN | TPR | TNR | PPV | ACC |
|---|---|---|---|---|---|---|---|---|
| BUSCA vs. HPA | 131 | 78 | 55 | 1130 | 0.70 | 0.94 | 0.63 | 0.90 |
| BUSCA vs. SubCell | 127 | 82 | 36 | 1149 | 0.78 | 0.93 | 0.61 | 0.92 |
| HPA vs. SubCell | 141 | 45 | 22 | 1186 | 0.87 | 0.96 | 0.76 | 0.95 |
| SubCell vs. HPA | 141 | 22 | 45 | 1186 | 0.76 | 0.98 | 0.87 | 0.95 |

TP – True positive predictions
FP – False positive predictions
FN – False negative predictions
TN – True negative predictions
TPR – True positive rate, sensitivity – TP / (TP + FN)
TNR – True negative rate, specificity – TN / (TN + FP)
PPV – Positive predictive value, precision – TP / (TP + FP)
ACC – Accuracy – (TP + TN) / (TP + TN + FP + FN)

The purpose of this analysis has been to assess the reliability of predicting mitochondrial localization with BUSCA, using HPA or SubCell as a reference. In addition, also the experimental data have been evaluated for reliability, using data from one method as “prediction” and the other as reference. The localization data have been simplified into “mitochondrial” and “everything else”. The comparisons have been done over the set of genes that has localization data in all three datasets, in total 1394 genes. This represents 42% of the genes with gene products analyzed in the KEGG pathways in the main paper.

The analysis shows that the specificity is very high, indicating that the predictions are quite reliable when a localization has been predicted to be mitochondrial.
